# The thalamic reuniens is associated with consolidation of non-spatial memory too

**DOI:** 10.1101/2023.04.01.535203

**Authors:** J. J. Hamilton, J. C. Dalrymple-Alford

## Abstract

The nucleus reuniens (RE) is situated in the midline thalamus and provides a key link between hippocampus and prefrontal cortex. This anatomical relationship positions the Re as an ideal candidate to facilitate memory consolidation. Evidence is, however, lacking whether this role extends beyond spatial memory and contextual fear memory, both of which are strongly associated with hippocampal function. We therefore trained intact male Long-Evans rats on an odour-trace-object paired-association task in which the explicit 10-second delay between paired items renders the task sensitive to hippocampal function. Neurons in the RE showed markedly increased immediate early gene (Zif268) activation when rats were re-tested for the previous non-spatial memory 25 days after acquisition training, relative to a group tested at 5-days post-acquisition, as well as a control group tested 25 days after acquisition but with a new pair of non-spatial stimuli, and home cage controls. The remote recall group also showed relatively augmented IEG expression in the superficial layers of the medial PFC (anterior cingulate cortex and prelimbic cortex). These findings support the conclusion that the RE is preferentially engaged during remote recall in this non-spatial task and thus has a role beyond spatial memory and contextual fear memory.

## 1. Introduction

The nucleus reuniens of the thalamus (RE) is ideally situated to mediate information exchange between the hippocampus and prefrontal cortex (PFC) and participate in a variety of higher cognitive functions (reviews: Dolleman-van der Weel et al., 2019; Ferraris et al., 2021; Mathiasen et al., 2020). One of the most interesting lines of research suggests that the RE is critical for long-term memory consolidation (Ferraris et al., 2021). Based on lesion studies in rats, the RE markedly reduces accurate recall 25 days after acquisition, but with little or no effect on short-term retrieval one to five days later, for spatial memory and contextual fear memory (Klein et al., 2019; Loureiro et al., 2012; Quet, Majchrzak, et al., 2020).

Evidence from immediate early gene (IEG) expression also supports the involvement of RE in long-term consolidation. C-Fos in the RE and PFC, and not the hippocampus, shows a relatively large increase for remote retrieval of spatial memory in the water maze (Klein et al., 2019; Loureiro et al., 2012). Another study found that the long-term retrieval of remote contextual fear memory also increased IEG activity in the RE, PFC and amygdala (Silva et al., 2019), although the RE increase was not evident in a second study (Quet, Majchrzak, et al. (2020). Together with lesion evidence, these findings suggest that the RE facilitates a dialogue between the PFC and the hippocampal system to support long-term memory consolidation and dynamic changes from a primarily hippocampal-dependent process to one that increasingly engages PFC activity.

Despite a role for the RE in remote memory recall when spatial information is required or when context refers to the background of diffuse multimodal stimuli, the broader relevance of the RE for memory consolidation may be limited. Specifically, one lesion study reported that the RE was not critical for long-term social-olfactory memory (Quet, Cassel, et al., 2020). These authors suggested that the social transmission of food memory may be supported by regions outside the hippocampus and this may explain why the RE was not involved in consolidation in this non-spatial task. The question remains, then, whether the engagement of the RE for long-term consolidation is limited to spatial and context memory specifically or more general processes that are sensitive to hippocampal dysfunction. The last point connects with debate as to whether the hippocampal system is responsible primarily for the acquisition of spatial and context memory or more general cognitive and relational representations. One critical issue is the presence of an explicit temporal feature of the memory (Ranganath, 2019; Whittington et al., 2022). For example, learning an object-odor non-spatial tasks when the two stimuli are presented simultaneously is not susceptible to hippocampal lesions in rats (Gilbert & Kesner, 2002). By contrast, the inclusion of a 10-second delay between the presentation of an object and an odor renders the non-spatial association memory susceptible to CA1 hippocampal lesions (Kesner et al., 2005), perhaps through mechanisms that link event sequences (Manns & Eichenbaum, 2005). In support of this, we have shown that dorsal CA1 neurons express increased IEG Zif268 activity during a 5-day recall when the memory representation included the 10-second (trace) delay between an odor and its associated object (i.e. an odor-trace-object memory) compared to recall when the paired association memory was trained without the delay (Hamilton & Dalrymple-Alford, 2022). Here, we provide evidence that the RE shows increased IEG activation during remote, relative to recent, recall of a previously acquired non-spatial association that incorporates such a temporal lag.

## 2. Materials and methods

### 2.1. Animals and housing conditions

Male Long-Evans rats, aged 12 months, were group-housed in Makrolon cages (48×28×22cm) on a reversed 12-hour light-dark cycle. Rats were maintained at 85% *ad libitum* body weight with testing during the dark phase. All procedures were approved (University of Canterbury Animal Ethics Committee and complied with ARRIVE guidelines). Three groups were established by random assignment. Post-acquisition, one group was tested for recent retention (5-days post-acquisition; n=8), the second for remote retention (25-days post-acquisition; n=8), and the third on a new association trained only at 25-days post-acquisition (25-day New, n=6); training and testing for all groups used a 10-second delay between the presentation of an odor and a subsequent object. There was also a home cage control group (n=9).

### 2.2. Non-spatial paired-associate memory task

For details on the apparatus and procedures, see Hamilton and Dalrymple-Alford (2022). Training on simple odor and simple object discriminations was conducted first to familiarise rats with the general “go/no-go” procedures. Acquisition for an odor-trace-object paired-association was conducted in the same red Perspex runway (93×26×26cm high) with vertically removable doors (26×26cm; Figure 1). The odor (e.g. 20ul of lemon or clove; Essential Oils, New Zealand) was mixed with 5ml of sunflower oil, and placed on a sponge (6.5×6×8mm) surrounding a plastic cap at the centre of the door at the end of the second compartment (15cm; B in Figure 1a); the cap presented a 1mg piece of chocolate, available after the rat was released from the 15cm-long start compartment. After removal of the door with the odorised sponge, the rat was retained for a 10-second delay in the third compartment (15cm). It was then allowed to run to the end of the runway to interact with a lightweight and visually distinct object. The object’s base was hinged to a food well beneath so the rat could search for additional chocolate reward. The presence of the reward under the object depended on whether the specific odor-trace-object pairing was a “correct” association. Black curtains enclosed the test area to minimise spatial cues. The observer sat within the curtains, adjacent to the runway and operated the doors.

**Figure 1.**
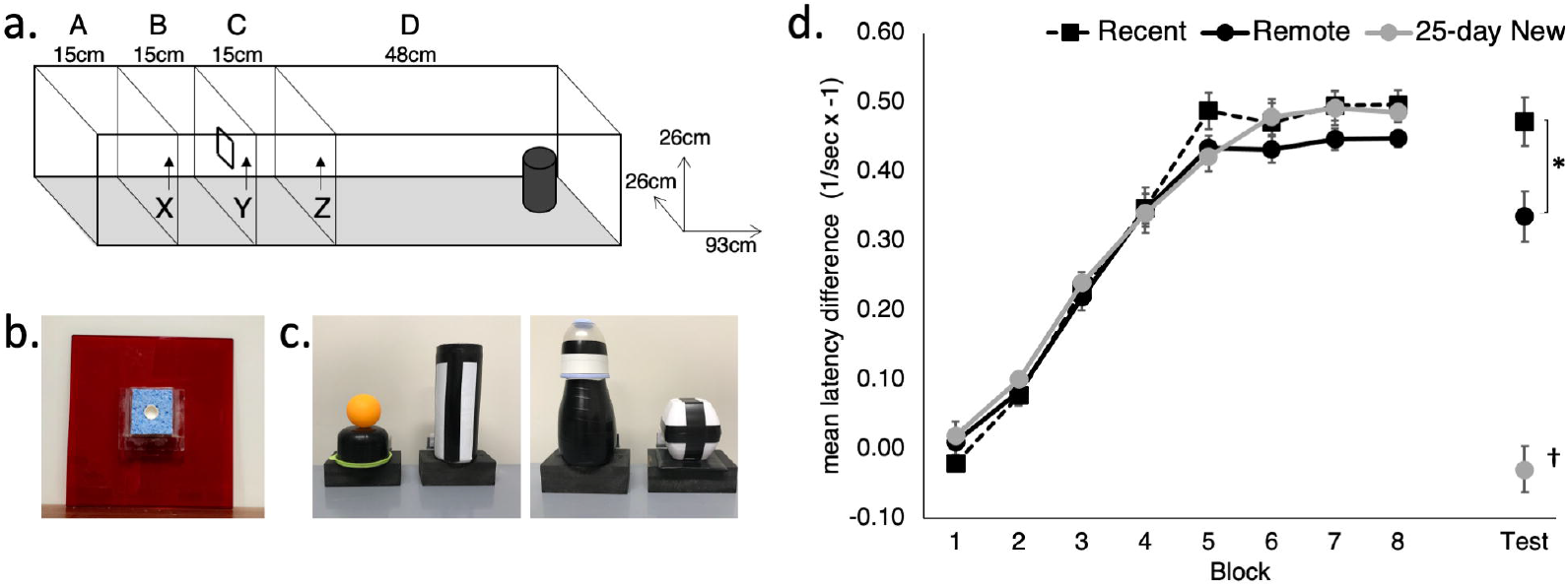
The odor-trace-object paired-associate memory task. **(a)** Rats started an odor-trace-object paired-associate trial in Compartment A and held there for 120 seconds on the first trial per day and 20 secs on the next eleven trials. Door X was removed to give access to the odor sponge **(b)** on door Y. On eating the food from the centre of the sponge, Door Y was removed and the rat was retained for a 10-second delay in Compartment C before Door Z was removed. Latency from removal of Door Z to interaction with the object **(c)** at the end of Compartment D was recorded. **(d)** Acquisition and retention of the odor-trace-object paired associate task in the three groups expressed as the latency difference between non-rewarded and rewarded pairings. The retention session was conducted 5 days (Recent) or 25 days (Remote; 25-day New) after the rat either reached criterion or the end of the acquisition period. The reciprocal latency (1/sec) was used because of the non-homogeneity of variance produced by raw latencies; reciprocal values have been inverted by multiplying by -1. Mean +/-standard errors. Recent, N = 8; Remote, N = 8; 25-day New, N = 6; * = significance at p<0.05; † = differs from the three other groups p<0.01.

Each rat received 12 massed daily trials, six go (rewarded) and six no-go (non-rewarded) in a pseudo-randomized order, with no more than three of either type run consecutively within session. Correct pairings for the paired-associate task were counterbalanced across rats. The critical measure was latency to push (nose or paw) the object after opening the last door. Rats learned that two odor-trace-object pairings were rewarded under the object (e.g. Odor 1 + Object A and Odor 2 + Object B) and that the two alternate pairings were not rewarded (i.e. Odor 1 + Object B and Odor 2 + Object A). A trial was ‘correct’ if the rat responded within 8 seconds on go trials or withheld responding for longer than 8s on no-go trials. Rats that reached a criterion of 80% correct trials over 3 consecutive days, after a minimum of 25 sessions (N = 15), were removed from further training and retention tested at either 5 days or 25 days later. There were 8 rats that continued for the maximum of 50 days of training, but all of these showed clear evidence of acquisition (3 rats each in Recent and Remote recall groups and 2 in the 25-day New group); their final latency difference between go and no-go trials was > 3.3 seconds with a mean of 3.8 seconds).

### 2.5. Post-acquisition retention test

Retention testing was conducted 5 days (recent recall) or 25 days (remote recall) after reaching criterion or else 50 days of training. The retention session used the same previously-learned association for the Recent and Remote groups. For the 25-day New learning group, a completely new odor-trace-object pairing was used per rat; this assessed the impact of a delayed return to testing conditions without explicit recall of the previously acquired association.

### 2.6. Histology

Immediately following the post-acquisition test (Recent, Remote, and 25-day New), or straight from an individual holding cage (Home Cage controls), rats were placed in a familiar dark, quiet room for 90 min prior to perfusion. Rats had experienced this procedure on two previous days. The rat was deeply anaesthetised with sodium pentobarbital (125 mg/ kg) and perfused transcardially with saline followed by 4% paraformaldehyde (PFA) in 0.1 M phosphate buffer (PB). Brains were post-fixed in 4% PFA and at least 48 h later placed in a long-term solution (20% glycerol in 0.1 M PB). Coronal 40-μm sections were collected using a freezing microtome (Thermo Fisher, UK) and stored in a cryo-protectant solution (30% glycerol, 30% ethylene glycol in 0.1 M PB) at -20°C prior to immunohistochemistry.

Free-floating sections were incubated in rabbit polyclonal Zif268 primary antibody (Egr-1; 1:1000 Cat# sc-110; Santa Cruz Biotechnology, USA) for 72 h at 4°C in PBSTx with 1% NGS, then incubated in biotinylated goat antirabbit secondary antibody (1:1000: Vector Laboratories BA-1000) overnight in PBSTx and 1% NGS. Following DAB visualisation (16-20 min, pending visual check) and mounting, Zif268-positive cell staining in each region of interest was photographed with a 10x objective using a light microscope (Leica, Germany). Automated cell counts were obtained through ImageJ analysis software (National Institute of Health, NIH, USA). Manual selections were made within the regions of interest per below and images converted to 8-bit grey scale, background was subtracted (rolling = 40), converted to mask and the watershed function applied. All neuronal cells above threshold (‘MaxEntropy’ threshold, circularity 0.65–1.0) were counted.

Regions of interest were selected for their potential involvement in memory recall. Based on the Paxinos and Watson (2014) atlas, we included two regions of the medial PFC with sections at +3.00 to +3.72 from Bregma. These were A32D (anterior cingulate region, ACC) and A32V (prelimbic region). We followed the descriptions provided by Vogt and Paxinos (2014) who delineated cell layers II, III and V in A32D, and layers II, III, V and VI in A32V. Layer delineations provided by Vogt and Watson (2014) were also followed for the retrosplenial cortex (−5.04 to -6.48 from Bregma) to identify layers in granular Rgb and granular Rga (both subregions, layers II, III, IV, V and VI; sections for Rga were centred at -6.36) and dysgranular Rdg (where layers II and III are deemed a single layer). For the midline thalamus, the RE and RH sections were taken between -1.80 and -2.16 from Bregma; the centromedial and paracentral intralaminar thalamic nuclei (ILt) were taken between -1.92 and -2.16 from Bregma. Sections for the dorsal and ventral hippocampal formation were based on atlas plates from -3.12 to -3.60 and -5.04 to -5.64 from Bregma, respectively.

Between two and six coronal sections containing a region of interest were selected. Within a cortical area with layers, such as A32D, the outermost layer of interest was identified and a middle portion of this was manually outlined to ensure clear separation from any adjacent region (e.g. within a dorsal to ventral aspect in A32D; transitions across a different number of layers in retrosplenial cortex subregions). Then subsequent layers were established beneath this used the landmarks and approximate depths described by Vogt and Paxinos (2014) for the prefrontal cortex and retrosplenial cortex subregions. For regions with no layers, as much of the region as possible within the photomicrograph was selected, while keeping separate of adjacent regions. Expression within each relevant layer or region of interest was quantified as the number of neurons per mm^2^ and the average Zif268-positive cell count derived across the relevant sections. Sections through the ILt were counted manually due to technical issues on the microscope.

### 2.7. Data analysis

ANOVA using Statistica (v13; Dell Inc.) evaluated mean differences across the groups (Recent, Remote, 25-day New, and Home Cage). Repeated measures factors were added for blocks of trials for the paired-associate memory task and Zif268 counts across related regions or subregions of interest. A reciprocal transformation of latency data for individual trials was used to establish homogeneity of variance. On each test day, the transformed latencies for individual trials generated a mean latency for the six rewarded trials and six non-rewarded trials with the difference used to evaluate performance. Latencies were carried forward for acquisition for rats that reached criterion. The final forty acquisition days were analysed, as running was more consistent after the first 10 days. To account for multiple comparisons and balance Type I and Type II errors, we used the significance level of p<0.02 for behavioral analyses and P<0.01 across regions of interest for Zif268 expression. Post hoc Newman–Keuls (N-K) tests assessed pairwise group differences. Simple main effects analysis was used when there were significant interactions involving repeated measures factors. Insufficient or missing brain sections in two rats reduced the degrees of freedom for some analyses; removal of these cases from other analyses did not change the findings. Effect sizes (Cohen’s d) were used to describe pairwise group differences in Zif268 expression in the RE.

## 3. Results

### 3.1. Behaviour

Rats in all groups rapidly acquired both the simple odor and simple object discrimination tasks at a similar rate (data not shown; simple odor discrimination, Group main effect, F(2,19)=0.08, p=0.92; simple object discrimination, F(2,19)=2.11, p=0.14).

The mean latency difference (based on reciprocal latencies) for paired-associate acquisition and performance on the retention test (Recent, Remote) and on the new odor-trace-object association (25-day New) is shown in Figure 1. An increasing latency difference was evident over training for the non-rewarded compared to the rewarded trials. All three groups acquired the odor-trace-object paired associate task at a similar rate (Figure 1; Group main effect, F(2,19)=0.52, p=0.60; Block main effect, F(7,14)=247.84, p<0.001; Group x Block interaction F(14,133)=0.71, p=0.75). The three groups did not differ on the final block of acquisition (Group at Block 10, F(2,19)=0.26, p=0.77). There was no correlation between mean latency difference in the final block of acquisition and the number of acquisition days (r(21)=-0.14, p>0.5).

Rats in the Recent and Remote recall groups showed clear retention of the acquired association compared to rats given the 25-day New odor-trace-object pairing (F(2,19)=9.63 p<0.001). Although the two recall groups exhibited similar responding on their last Block of acquisition (Block 10), the Recent recall group showed better retention than was shown by the Remote recall group (p<0.02).

### 3.2 Zif268

Figure 2 shows Zif268 expression in the RE, RH, and the medial prefrontal A32D (ACC) and A32V (prelimbic) regions of interest for the four groups (Recent, Remote, 25-day New, and Home Cage controls). In the RE, all three trained groups showed higher expression than was found in the Home Cage group (Group main effect F(3,25)=64.72, p<0.001), suggesting that engagement in and general memory for the behavioral task *per se* associates with increased activity in the RE. Critically, N-K post hoc analysis confirmed that all four groups differed from each other for Zif268 expression in the RE (p<0.004). Remote recall of the odor-trace-object paired-associate memory produced markedly higher Zif268 expression in the RE compared to home cage group (by a factor of 298%; effect size d = 9.91), recent recall (161% increase; d = 4.13) and exposure to a new association at 25-days (120% increase; d = 1.87). To a lesser extent, both recent recall and exposure to the new association also produced greater levels of expression compared to Home Cage (Recent: 185% increase; d = 4.25; 25-day New: 247% increase; d = 7.35). There was no correlation (r(19)=0.21, p>0.3) between the mean latency difference score on the retention test and zif268 expression in the RE within the individual groups: Remote, r(6)=0.24, p>0.6; Recent, r(7)=-0.60, p>0.1; 25-day New, r(5)=-0.11, p>0.8). The days taken to reach criterion was also not associated with Zif268 expression in the RE (r(19)=-0.31, p>0.17).

**Figure 2.**
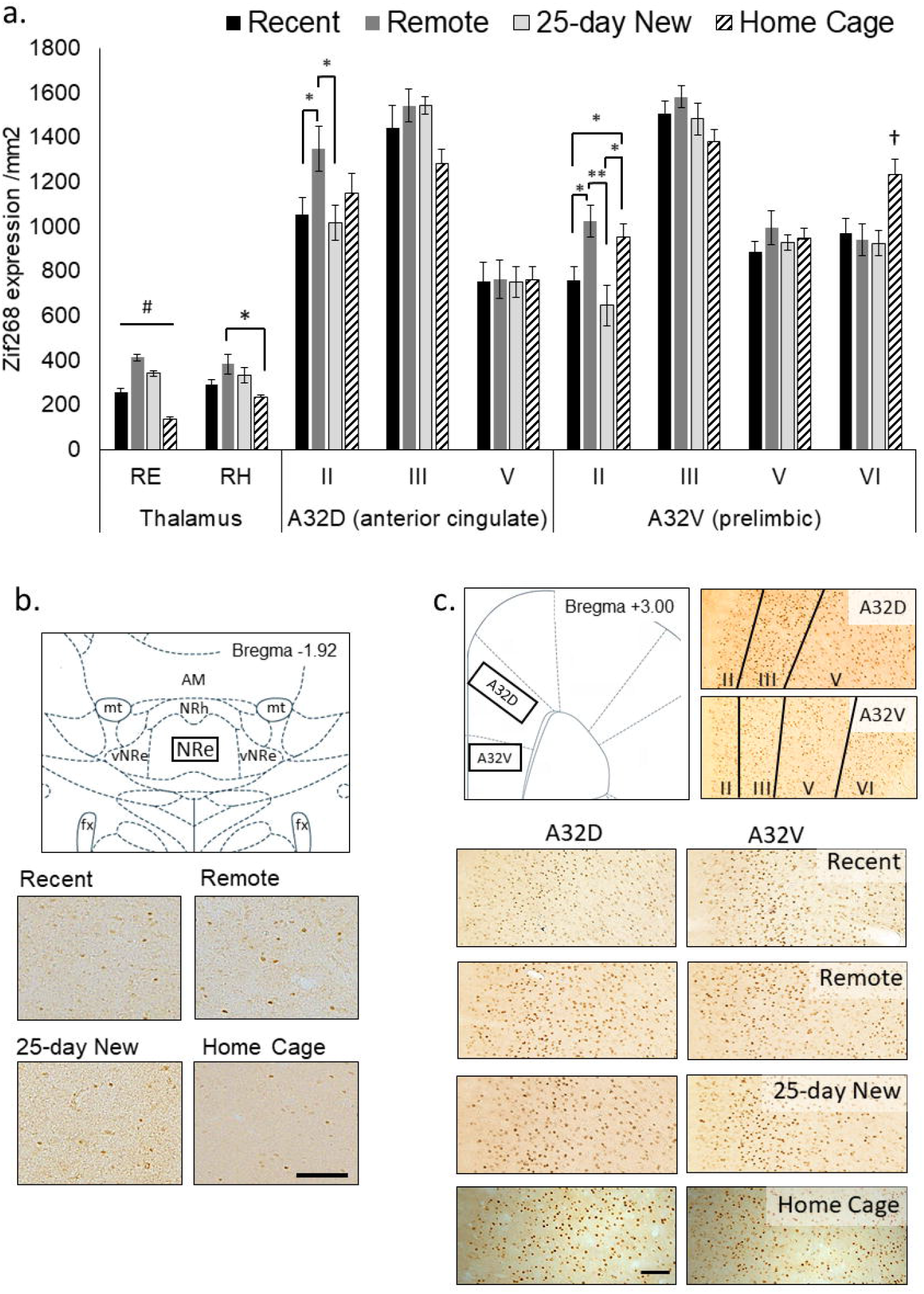
**(a)** Zif268 expression per mm^2^ (Mean +/-standard errors) assessed in the nucleus reuniens (RE), rhomboid nucleus (RH), and prefrontal cortex regions (A32D, anterior cingulate, Layers II, III and V; A32V, prelimbic, Layers II, III, V and VI) in the three groups immediately after testing at 5-days (Recent) or 25 days (Remote; 25-day New) after acquisition, as well as Home Cage controls. **(b)** Schematic diagram showing the RE region assessed and expression in 10x magnification photomicrographs for each group. The RE was assessed from -1.80 to -2.16 from Bregma. **(c)** Schematic diagram of the mPFC (A32D and A32V; +3.00 to +3.72 from Bregma) regions and the corresponding layers assessed shown in 10x magnification photomicrographs, and expression in photomicrographs for each group. The Home Cage control group was processed separately, but using the same Zif268 primary antibody batch number. Variation in background staining was accommodated by adjusting the background threshold in ImageJ. Recent N = 8; Remote N = 8; 25-day New N = 6; Home Cage N = 9; Black bar = 100μm; * = significance at p <0.05; ** = p<0.01; # = all groups differ to each other at p<0.01; † = differs from the three other groups p<0.01.

A similar, albeit weaker, pattern of Zif268 expression to that of the RE was apparent in the RH (Group main effect, F(3,25)=4.42, p=0.01). However, post-hoc N-K revealed a significant difference only between Remote and Home Cage groups (p=0.01) with Recent and 25-day New producing non-significantly lower levels of expression compared to the Remote group (p<0.10).

There was no Group main effect for the A32D (ACC) region, aggregated across all three layers (F(3,27)=1.18, p=0.35). A Group x Layer interaction (F(6,54)=3.49, p=0.005) was, however, evident with the Remote recall group showing higher Zif268 expression in the superficial Layer II than in both the Recent recall group (F(1,27)=5.62, p=0.02) and the 25-day New learning group (F(1,27)=6.16, p=0.01). Zif268 expression in Home Cage group was not significantly different to any other group (p<0.12). There were no group differences for the deeper layers.

Across the cell layers of the A32V (prelimbic) region of the medial PFC, Zif268 expression showed a similar pattern to that seen in A32D. That is, there was no Group main effect (F(3,27)=1.95,p=0.14) but a Group x Layer interaction (F(9,81)=7.13, p<0.001). The interaction was again driven by elevated expression in superficial Layer II in the Remote recall group, with expression in this group significantly higher compared to both Recent (F(1,27)=7.66, p=0.01) and 25-day New groups (F(1,27)=13.56, p<.002). The Home Cage group showed an intermediate level, but significantly higher than both Recent and 25-day New groups (p<0.05). In the deeper Layer VI, however, the Home Cage group showed elevated Zif268 compared to all three trained groups (p<0.01), but there were no differences in the intermediate layers (Layers III and V).

Figure 3 shows Zif268 expression in the hippocampus, ILt and retrosplenial cortex in the four groups. Zif268 expression across the dorsal and ventral CA1 did not differ between the trained groups or the Home Cage group (Group main effect, F(3,27)=1.39, p=0.26). There was also no Group by Dorsal/Ventral interaction (F(3,27)=1.64, p=0.19). Across both the dorsal and ventral CA3, however, Home Cage rats expressed much higher Zif268 levels (Group main effect, F(3,26)=44.12, p<0.001), while the three trained groups did not differ from each other (N-K p>0.1).There was no Group by Dorsal/Ventral interaction for CA3 (F(3,26)=1.44, p=0.25). In the ILt, rats in the Home Cage group expressed less Zif268 than all three trained groups (Group main effect, F(3,25)=5.36, p=0.005), but the trained groups did not differ from each other (p>0.5). There was no Group by ILt sub-region interaction (F(3,25)=1.20, p=0.32).

**Figure 3.**
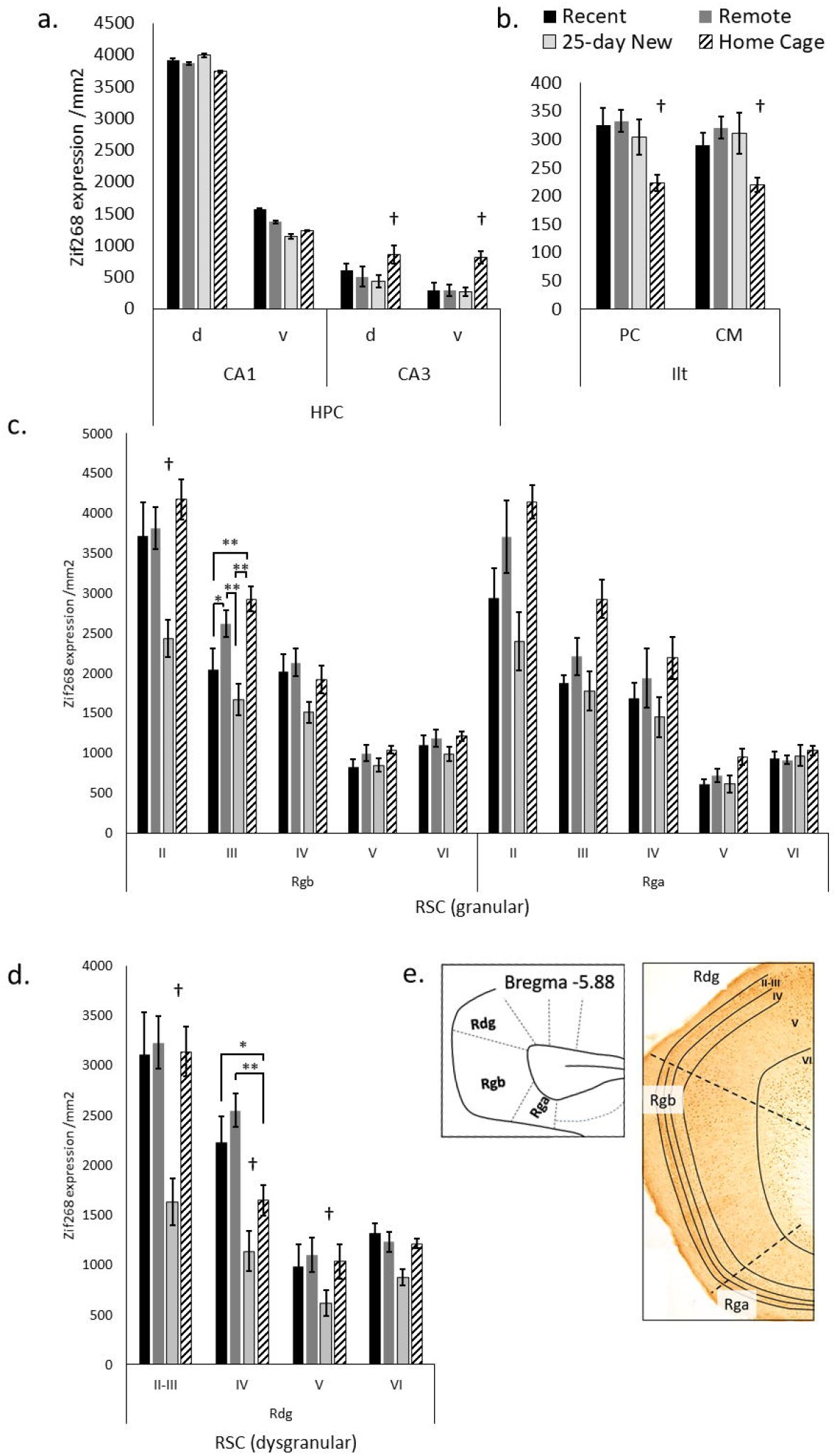
Zif268 expression per mm^2^ (Mean +/-standard errors) assessed in the **(a)** dorsal (d) and ventral (v) hippocampal CA1 and CA3, **(b)** paracentral (PC) and centromedial (CM) intralaminar thalamic nuclei (ILt), and **(c)** the retrosplenial cortex regions: granular Rgb, granular Rga, and **(d)** dysgranular Rdg in the three trained groups and the Home Cage group. **(e)** Schematic diagram of the caudal regions of the retrosplenial cortex with examples of layers II-VI shown in the photomicrograph. Note, Rga was quantified only in the posterior retrosplenial sections (−5.64 to -6.48 from Bregma), while Rgb and Rdg had sections compared from both anterior and posterior retrosplenial cortex (−5.04 to -6.48 from Bregma). The deeper increase in thickness towards to more posterior aspects of the Rgb and Rdg. Recent N = 8; Remote N = 8; 25-day New N = 6; Home Cage N = 9; * = significance at p <0.05; ** = p<0.01; † = differs from the three other groups p<0.01.

In the Rgb subregion of the retrosplenial cortex, the 25-day New group (that is, with a new odor-trace-object pairing in the retention test) showed lower Zif268 expression compared to the other groups (Group main effect, F(3,27)=6.28, p=0.002). This difference was larger relative to the Home Cage group (N-K, p<0.001) than the Recent (p=0.02) and Remote (p=0.003) groups. The group differences, however, also varied across Rgb layers (Group x Layer interaction, F(12,108)=3.65, p<0.001), diminishing across depth of layer. In both Layer II and III, the 25-day New group were different to the other three groups; in layer III, the Remote Group now also showed increased levels compared to the Recent Group. There were no group differences for layers V and VI (F < 1.61).

For the Rga subregion, there was also a Group main effect (F(3,25)=7.44, p=0.001) with a graded decrease in Zif268 expression from Home Cage, to Remote Group, Recent Group and 25-day New group. The Home Cage group showed significantly higher expression than both Recent (N-K, p=0.006) and 25-day New groups (N-K, p=0.001), but no other differences reached significance. Although these differences reduced with increase in depth of layer, the Group by Layer interaction was not significant (F(12,100)=1.81, p<0.06).

In the Rdg subregion, the 25-day New group again expressed lower levels of Zif268 than the other three groups (Group main effect, F(3,27)=19.09, p<0.001; pairwise N-K, p<0.001). The Recent, Remote and Home Cage groups did do not differ (N-K p>0.1). There was also a Group by Layer interaction (F(9,81)=5.91, p<0.001) due to the differences with the 25-day New Group diminishing across deeper layers (p<0.005 except layer VI p=0.07). In Layer IV, the Home Cage group expressed an intermediate level of Zif268, which was significantly different from the Recent (p=0.02) and Remote (p<0.001) groups. No other differences reached significance.

## 4. Discussion

The current study builds on evidence that the RE makes an important contribution to consolidation and remote recall for spatial memory (Klein et al., 2019; Loureiro et al., 2012) and contextual fear memory (Quet, Majchrzak, et al., 2020). We found a robust increase in RE activity during long-term, but not short-term, retrieval of a non-spatial associative memory in which the inclusion of a brief (10-second trace) between the presentation of the odor and the rat’s interaction with the object renders this task hippocampal-dependent (Kesner et al., 2005). The current study found a large effect size for Zif268 expression in the RE for remote recall relative to when retention was tested 5 days after acquisition, and compared to two control conditions (25-day retesting but with a new association; and home cage control). There was only weak evidence that the RH was engaged in a similar fashion for remote recall, although a broadly similar pattern of IEG activation to that of RE expression was found across the three trained groups. The value of this paired-associate task is that it offers the opportunity to explore memory for an association between arbitrary non-spatial stimuli, the impairment of which is a core feature of the amnesic syndrome (Turriziani et al., 2004).

The engagement of the RE may also be relevant to performance of the odor-trace-object paired-associate task, because all three trained groups showed a marked increase in Zif268 at the time of testing compared to home cage controls. In addition, all three trained groups showed increased Zif268 activation in both paracentral and centromedial sub-regions of the ILt compared to home cage controls. By contrast, inhibition of the superficial layer of the retrosplenial cortex may be relevant to new learning in this task as there was a reduced IEG activity in the 25-day New group compared to the two recall groups and the home cage controls.

Increased IEG activity in the superficial layer in the medial PFC regions with remote recall in our non-spatial task, using Zif268 as a neuronal marker, is also consistent with previous work on memory consolidation. Remote recall in a spatial water maze task was accompanied by increased c-Fos expression in the prelimbic cortex and ACC, and with increased spinogenesis in superficial layers; these effects were attenuated by RE lesions (Klein et al., 2019). Together, the findings suggest that communication between the RE, the medial prefrontal cortex and the hippocampus may be relevant for remote retrieval beyond spatial and contextual memory. This broadens the role of the RE in hippocampal-diencephalic-cortical connectivity for consolidation and memory persistence across different kinds of memory tasks.

We selected Zif268 because this IEG marker provided prior evidence of engagement by dorsal CA1 neurons for 5-day recall after training with a 10-second trace between the odor and object stimuli compared to 5-day retention after training without the inter-item delay (Hamilton & Dalrymple-Alford, 2022). Previous research on IEG activation in the RE has used c-Fos as the functional marker (e.g., Klein et al., 2019; Loureiro et al., 2012). However, similarities rather than differences can usually be expected across the two functional markers and in any case our results were positive with Zif268. For example, direct comparison of Zif268 and c-Fos in the cortex following remote recall compared to recent recall of a five-arm spatial reference memory produced equivalent effects (Maviel et al., 2004). Thus, either IEG marker seems suitable for functional brain imaging associated with memory consolidation.

A previous study suggested that the RE may not be critical for long-term consolidation of non-spatial memory (Quet, Cassel, et al., 2020). However, there are differences in task characteristics between this previous study and those used in the current study. Quet, Cassel, et al. (2020) determined that pre-acquisition lesions of the RE did not affect retrieval of social-olfactory memory, and commented on mixed evidence for the role of the hippocampus in that task as an explanation for this lack of effect. By contrast, the current study used very different task procedures and focused on neuronal activity at the time of recall in a specific odor-trace-object task that other work suggests preferentially engages dorsal hippocampal CA1 neurons (Hamilton & Dalrymple-Alford, 2022; Kesner et al., 2005). Moreover, the current study was focused on the presence and pattern of neuronal activation associated with remote recall and was not designed to test the criticality of the RE per se at remote retrieval.

While RE activity at the time of recall associates with remote retrieval of contextual memory, spatial memory, and now non-spatial odor-trace-object association memory, there appear to be multiple thalamic sites that could support remote retrieval in the absence of RE activity at the time of recall. There is evidence that disengagement of the anterodorsal thalamic nuclei facilitates remote contextual fear memory recall (Vetere et al., 2021), whereas activation of the anteromedial thalamic nuclei and ILt are relevant for remote memory of differentially-reinforced (salient) contexts and spatial cues, respectively (Lopez et al., 2009; Toader et al., 2023). Unfortunately, we had minimal Zif268 staining in the anterior thalamic nuclei in our material, so we cannot address their involvement in our study. Nonetheless, alternate thalamic sites may reduce the critical importance of the RE during either systems level consolidation or remote recall. If several nuclei contribute to consolidation and recall of a remotely acquired memory, this could explain evidence that environmental enrichment reinstated remote recall and elevated medial PFC activation in rats with RE lesions (Ali et al., 2017). Similarly, the failure of RE lesions to impair remote recall of social-olfactory memory may be due to the relative importance of alternate thalamic structures for that memory (Quet, Cassel, et al., 2020). Future work is needed to determine how multiple neural circuits that engage the midline thalamus cooperate in the process of acquisition, consolidation, and long-term memory retrieval for different types of events.

## Data availability statement

Data will be available on request.

## Author contributions

JJH and JCDA: funding, concept and design, writing and editing drafts, statistical analysis and interpretation; JJH: conducting experiment.

## Acknowledgments

This research was supported by University of Canterbury equipment and research grants and Early Career support (JJH) from Brain Research New Zealand – Rangahau Roro Aotearoa and Royal Society of New Zealand, Canterbury Branch.

